# Locus Coeruleus α-Synuclein Overexpression Induces Prodromal Parkinsonian Features in Mice

**DOI:** 10.1101/2025.11.15.688542

**Authors:** Jone Razquin, Laura De Las Heras-García, Andrea Vaquero-Rodríguez, Gloria González-Aseguinolaza, Edgar Soria-Gómez, Jérôme Baufreton, Harkaitz Bengoetxea, Naiara Ortuzar, Jorge E. Ortega, Cristina Miguélez

**Author notes:** Correspondence to: Cristina Miguélez, Department of Pharmacology, Faculty of Medicine and Nursing, University of the Basque Country UPV/EHU, Barrio Sarriena s/n 48940 Leioa, Spain.

## Abstract

The locus coeruleus is one of the first brain regions to develop α-synuclein pathology during the prodromal phase of Parkinson’s Disease, contributing significantly to non-motor symptoms such as cognitive decline, mood alterations or sleep disturbances. However, the precise role of the locus coeruleus in the early stages of the disease remains unclear.

To address this, we developed and characterized a novel mouse model based on the PRSx8-driven overexpression of human α-synuclein in noradrenergic neurons of the locus coeruleus using an adeno-associated viral vector. Animals were assessed at 1 and 3 months post-injection using an integrated battery of histological, behavioural, neurochemical, and electrophysiological analyses.

We observed robust accumulation of phosphorylated α-synuclein in the locus coeruleus, along with widespread propagation to projection areas, including the hippocampus, prefrontal cortex, and dorsal raphe. Despite the absence of neuronal loss in the locus coeruleus, we identified reduced noradrenergic axonal integrity and marked decreases in both noradrenaline and serotonin levels, highlighting a disruption of neurochemical balance at an early stage. The electrophysiological data revealed transient alterations in the excitability and intrinsic properties of locus coeruleus neurons in a sex-dependent manner, suggesting that α-synuclein pathology induces early functional disruption prior to overt neurodegeneration. Behavioural outcomes demonstrated selective cognitive deficits and mild anxiety-like behaviours, while other non-motor functions remained preserved. Importantly, although several pathological and functional alterations were evident in the initial phases, behavioural impairments were also maintained at later time points, indicating that locus coeruleus-driven pathology exerts long-lasting consequences. Interestingly, female mice exhibited colon shortening, providing evidence of a sex-specific pathological process at the level of the gut-brain axis.

We believe that this model accurately replicates key prodromal features of Parkinson’s disease, demonstrating that α-synuclein-induced pathology in the locus coeruleus leads to functional circuit impairment and cognitive decline. Our findings emphasize the locus coeruleus as a critical site of early vulnerability in Parkinson’s Disease and underscore the importance of sex as a biological variable influencing disease progression and therapeutic response.

## Introduction

Parkinson’s disease (PD) is the most prevalent movement disorder and the second most common neurodegenerative disease worldwide, affecting approximately 0.1% of the global population.^1^ Its pathological hallmark is the accumulation of α-synuclein (a-syn) aggregates, forming Lewy bodies, which compromise motor nuclei such as the substantia nigra pars compacta (SNc), resulting in the characteristic motor symptoms of bradykinesia, tremor, and rigidity. However, Lewy body pathology also develops in non-motor regions up to two decades before motor symptoms emerge, particularly in the locus coeruleus (LC), one of the earliest brainstem structures to be affected.^2,3^

*Post-mortem* studies consistently demonstrate significant LC neuronal loss and a-syn pathology, with LC degeneration occurring before that of dopaminergic neurons in the SNc.^4–7^ Moreover, neuromelanin-sensitive MRI studies reveal progressive loss of LC signal in both genetic and idiopathic PD.^8–10^ Noradrenergic dysfunction affects a broad range of projection targets, including the prefrontal cortex (PFC), motor cortices, striatum, thalamus, hypothalamus, and cerebellum,^11–16^ as well as the peripheral autonomic system.^17,18^ This disruption is reflected in reduced cerebrospinal fluid levels of the noradrenaline (NA) metabolite dihydroxyphenylglycol^19^ and decreased dopamine-β-hydroxylase (DBH) activity.^20^

Noradrenergic impairment has been linked to several non-motor symptoms of PD, including autonomic dysfunction, depression, anxiety, and cognitive decline.^16,21–25^ Orthostatic hypotension, urinary, and sexual dysfunction are correlated with reduced LC integrity and lower NA levels.^26–29^ Similarly, LC degeneration is strongly associated with neuropsychiatric disturbances. Depressed PD patients exhibit greater noradrenergic deficits than non-depressed patients. This has been observed through lower contrast-to-noise ratios, neuronal loss and gliosis in the LC^30,31^ along with reduced NA and DA innervation of limbic areas.^14^ Imaging studies have also demonstrated that LC degeneration is associated with higher apathy scores^32,33^ and cognitive impairment,^16,32,34^ particularly affecting attention and working memory domains.^35^

Experimental studies also highlight the LC as a critical modulator of disease progression. In PD models, NA depletion exacerbates nigrostriatal degeneration^36–38^ whereas enhancing noradrenergic tone promotes SNc neuronal survival.^39–41^ Preclinical approaches using combined NA/DA lesions or genetic models provide additional evidence for a key role of LC dysfunction in the emergence of non-motor PD features.^42–44^ These findings closely mirror the deficits observed in patients.

Together, clinical and preclinical data underscore a crucial role for noradrenergic dysfunction in PD pathogenesis. However, the contribution of a-syn accumulation specifically within the LC to the early features of the disease remains unclear. Here, we investigate the impact of adeno-associated virus (AAV)-mediated selective human a-syn overexpression in LC neurons, assessing its contribution to noradrenergic dysfunction and the emergence of prodromal PD symptoms.

## Material and methods

A detailed description of the experimental procedures is provided in the Supplementary Material.

### Animals

Male and female C57BL/6J mice (Envigo, Spain), 8 weeks old at the start of the experiment, were used in this study. Animals were housed in groups of six under standard laboratory conditions, with *ad libitum* access to food and water.

### Experimental design and viral vector injection

AAV9 vectors were generated carrying either GFP, human α-synuclein (hα-syn), or no transgene, all driven by the PRSx8 promoter and containing the WPRE element: AAV9-PRSx8-GFP-WPRE, and AAV9-PRSx8-hasynWT-WPRE and AAV9-PRSx8-WPRE. Behavioural, histological, neurochemical and electrophysiological experiments were performed 1 and 3 months after stereotaxic injection of the viral vectors either bilaterally or unilaterally in the LC (Fig. 1). Sample sizes were determined based on our previous similar studies. For behavioural experiments the order of testing was randomized using Excel’s RAND() function (one random number per animal). The experimenter was blinded to group allocation during data collection and analysis.

**Figure 1.**
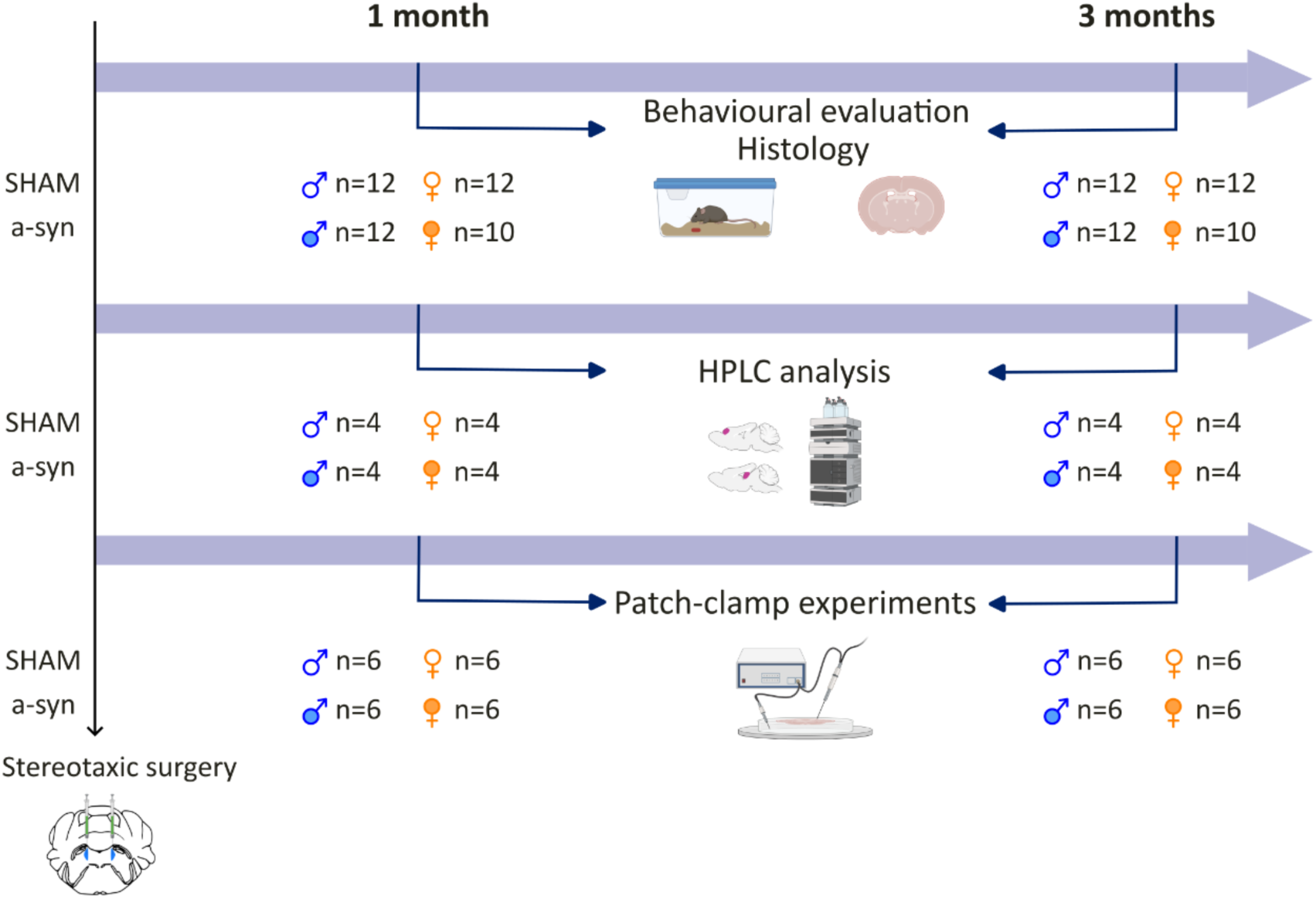
Experimental design. Stereotaxic surgery was performed in male and female mice to inject the viral vector. After establishing the a-syn model, behavioural tests, histological analyses, HPLC measurements, and in vitro electrophysiological recordings were conducted at 1 and 3 months post-surgery.

### Immunohistochemistry and analysis

Immunofluorescence procedures were performed to assess a-syn overexpression and its co-expression with noradrenergic neurons (tyrosine hydroxylase, TH) or projections (noradrenergic transporter, NET). Additional assays were conducted to detect GFP and phosphorylated a-syn (a-synP). DBH axon expression in the hippocampus (HPC) and the identification of in vitro recorded neurons using streptavidin were also assessed. LC neurons were quantified based on TH immunofluorescence (TH+ neurons/mm²), and vector transduction efficiency was assessed using TH and a-syn double immunofluorescence. DAB immunostaining was used to detect TH expression in SNc slices and a-syn across the brain axis. Stereological quantification was used for SNc neuron counts and optical density for a-syn expression.

### Brain Tissue NA and 5-HT Analysis

Brains were dissected and the PFC, the remaining cortical area and HPC were isolated, frozen, and processed for high-performance liquid chromatography (HPLC) with electrochemical detection (VT-03, Antec Scientific) at 0.3V. Chromatographic data were analysed using Chemstation plus software, and monoamine concentrations were determined via linear regression of standard curves (0.5–400 nM). The caudal brain was used for immunohistochemical verifications of a-syn overexpression.

### Spleen weight and colon measurement

The spleen and colon were removed for measurement and weighing after sacrificing the animal, when perfusion was not performed.

### Patch-clamp electrophysiology

Mice were anesthetized with 5% isoflurane, decapitated, and brains rapidly removed. Coronal LC slices (220 μm) were prepared using a vibratome (Microm HM 650V) and incubated in artificial cerebrospinal fluid (ACSF) at 35°C. Slices were perfused with oxygenated ACSF (32–34°C) during recordings. LC neurons were visualized with a microscope (Nikon Eclipse E600FN, 60× water-immersion objective). Whole-cell recordings were obtained using borosilicate pipettes (3–6 MΩ) filled with internal solution containing K-Gluconate and Neurobiotin (1%). Signals were recorded using an Axopatch 200B amplifier and Digidata 1322A digitizer (Molecular Devices) controlled by Clampex 10.3. Voltage-clamp recordings were low-pass filtered at 2 kHz and sampled at 10 kHz, whereas current-clamp recordings were filtered at 5 kHz and sampled at 20 kHz. Junction potential was 15 mV and was not corrected. Spontaneous firing rates were recorded in the cell-attached configuration with neurons held at −60 mV. LC neurons were identified by a resting inwardly rectifying potassium conductance, measured by stepping the membrane potential from −40 to −120 mV in −10 mV steps. Passive membrane properties were assessed by injecting −5 mV steps to measure capacitance and resistance. In current-clamp mode, current pulses (−300 to +300 pA in 25 pA steps) were injected to assess evoked firing rates and generate current-voltage curves. Action potential (AP) properties were analysed from the first sweep without current injection. Data were analysed off-line using Clampfit 10.7 software (Molecular Devices).

### Behavioural assessment

#### Open field test

Locomotion was assessed in an open field (OF) arena (44 × 44 × 35 cm) using Actitrack software (Panlab, Spain). Movement was detected using infrared beams, and the arena was divided into central and peripheral zones to assess anxiety-like behaviour. Mice were placed in the centre, and activity was recorded for 10 min.^45^

#### Sucrose preference test

Anhedonia was assessed using the sucrose consumption test, a non-invasive measure of depressive-like behaviour. Mice were habituated to two bottles to prevent side bias. During testing, one bottle contained water and the other a 3% sucrose solution, with randomized placement. Consumption was measured by weighing bottles at 2 and 24 h.

#### Buried food task

Olfactory impairment was assessed using the buried food task. Mice were familiarized with a scented pellet (Versele-Laga’s Complete Crock®, cheese or apple flavours) for 72 h. To prevent bias, different flavours were used at 1 and 3 month tests. On day 1, mice explored a sawdust-filled cage (3 cm deep) for 5 min. On day 2, the pellet was buried, and mice were given 5 min to locate it. No food deprivation was applied prior to testing.^46^

#### Hot plate

Hyperalgesia was assessed using the hot plate test. Mice were placed on a 52.5°C metallic surface (Digital DS-37 Socrel, Milan, Italy) surrounded by a 40 cm plexiglass wall. Pain latency (paw licking or jumping) was recorded, with a 45 s time limit to prevent tissue damage.^47^

#### Novel object recognition test

Cognitive ability was assessed using the novel object recognition (NOR) test in an L-shaped maze (8 × 37 × 14.5 cm).^48^ The test lasted 9 min per day for three consecutive days. Exploration (nose ≤2 cm from object) was scored using the Behaviour Scoring Panel. Heat maps were generated with EthoVision® (Noldus, Netherlands). The discrimination index was calculated as (Tnew−Told) / (Tnew+Told).

### Statistical analysis

Data were analysed using GraphPad Prism v8.0.1 (GraphPad Software Inc., USA). Results are expressed as mean ± standard error of the mean (SEM). Single-variable comparisons were assessed using unpaired Student’s t-tests. Two-way ANOVA was used to evaluate the effects of group (sham vs a-syn) and time. Tukey’s post hoc test was applied when significant interactions were detected. Statistical significance was defined as P < 0.05.

### Data availability

Data are available on request to the corresponding author.

## Results

### Targeted a-syn overexpression in LC neurons and propagation along noradrenergic projections

To evaluate the efficiency of AAV-PRSx8-hsyn mediated overexpressing a-syn immunofluorescence analyses were conducted in brain tissue collected at 1 and 3 months post-injection. A-syn was selectively expressed in TH+ neurons of the LC at both time points. High-magnification images confirmed its localization within noradrenergic neurons (Fig. 2A). Quantitative analysis demonstrated high transduction efficiency, with 78.38% and 70.95% of TH+ neurons expressing a-syn at 1-and 3-months respectively. Consistently, GFP expression driven by the same PRSx8 promoter was selectively observed in LC neurons (Supplementary Fig. 1).

**Figure 2.**
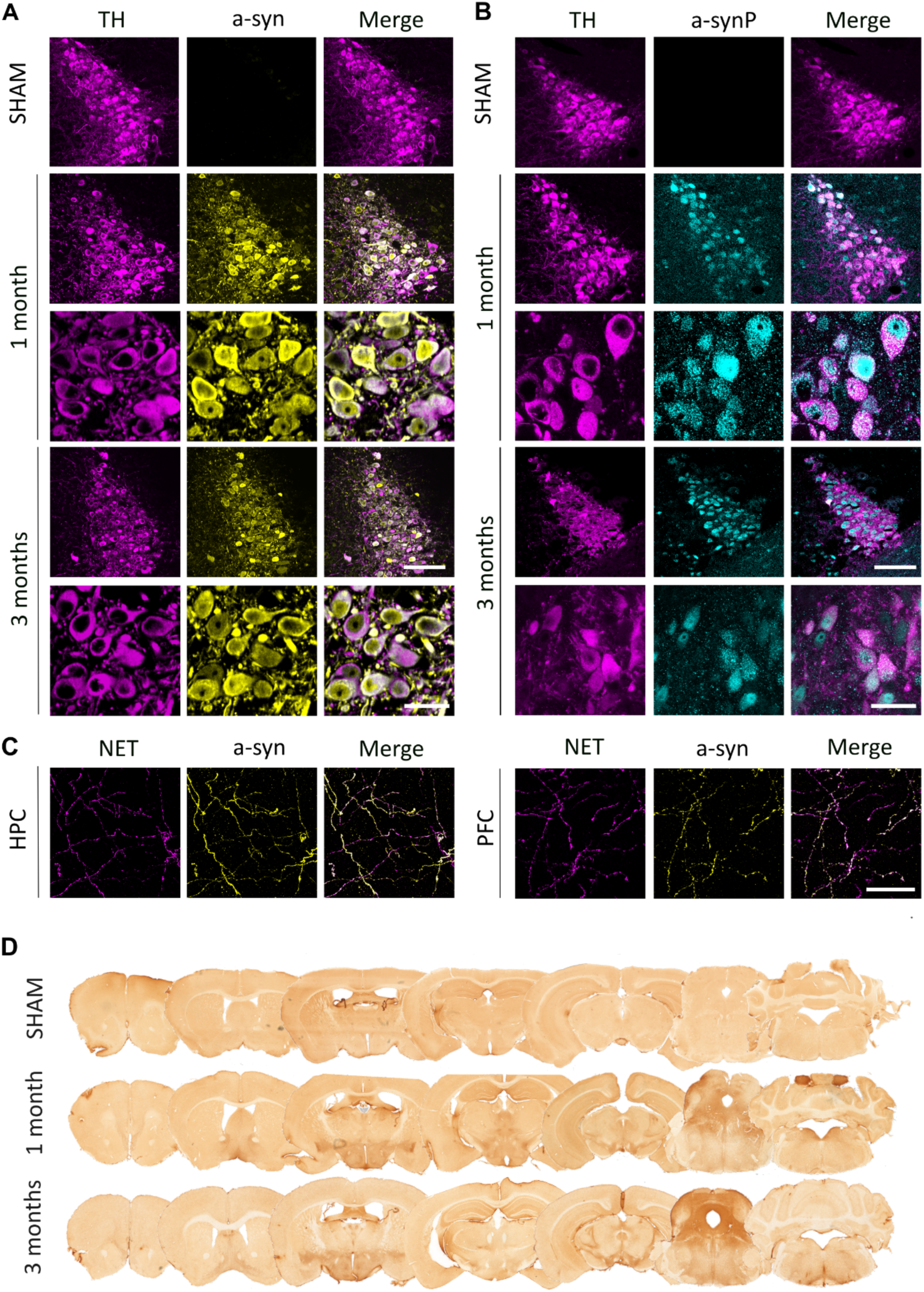
A-syn expression in the Locus Coeruleus. (**A**) Representative images of TH+ (magenta) and a-syn (yellow) colocalization in the LC at 1 and 3 months post-surgery. (**B**) Phosphorylated a-syn (cyan) remains restricted to LC neuronal somas. Scale bars: 100 µm (20×), 25 µm (63×). (**C**) Colocalization of a-syn with NET in the hippocampus and prefrontal cortex. Scale bars: 25 µm (63x). (**D**) A-syn distribution along the rostro-caudal axis in bilaterally injected brains.

The presence of a-synP, a key pathological hallmark of protein aggregation was also assessed.^49^ While monomeric a-syn was localized in the LC and distributed throughout noradrenergic neuronal projections, a-synP was predominantly confined to LC somas, at both 1 and 3 months (Fig. 2B). Altogether, these findings demonstrate that AAV-PRSx8-ha-syn drives selective and sustained a-syn expression in locus coeruleus noradrenergic neurons, leading to its phosphorylation and providing a robust model to investigate early pathological events associated with a-syn accumulation in this nucleus.

To determine whether a-syn overexpression propagates from LC neuronal somas along noradrenergic axons, double immunofluorescence staining was performed using antibodies against NET and a-syn. LC projection areas, including the PFC and the CA1 layer of the HPC, exhibited a fibre-like distribution of a-syn staining. These a-syn+ fibres clearly colocalized with NET-labelled LC axons (Fig. 2C). These results indicate that a-syn is localized within noradrenergic fibres originating from the LC, supporting the hypothesis that it spreads primarily via synaptic transmission along LC projections.

To analyse a-syn spreading across LC-projecting areas, a-syn immunohistochemistry was performed on serial brain sections encompassing the entire brain. At 1 month, a-syn+ fibres were already abundant in LC projection areas associated with PD, such as the SNc, subthalamic nucleus, PFC, HPC, motor cortex, and dorsal raphe (DR) (Fig. 2D). To further analyse a-syn propagation patterns and interhemispheric spread, 14 male mice received unilateral injections of the viral vector. Brains were analysed at 1 and 3 months post-injection, and both qualitative and quantitative analyses were performed. Regarding interhemispheric propagation, while most regions exhibited reduced a-syn+ fibre density on the contralateral side of the injection, certain nuclei such as the HPC and ventral tegmental area showed symmetrical distribution of a-syn fibres across both hemispheres. Notably, the LC contralateral to the injection site displayed less a-syn expression compared with more distal areas, including the DR, SNc, subthalamic nucleus and HPC (Fig. 3A and Supplementary Table 2). Semiquantitative analysis confirmed that the HPC and DR exhibited the highest densities of a-syn expression. By 3 months, a slight increase in a-syn optical density was observed, mainly in the HPC (Fig. 3B). Despite robust LC overexpression, interindividual variability in a-syn levels across different brain regions was evident.

**Figure 3.**
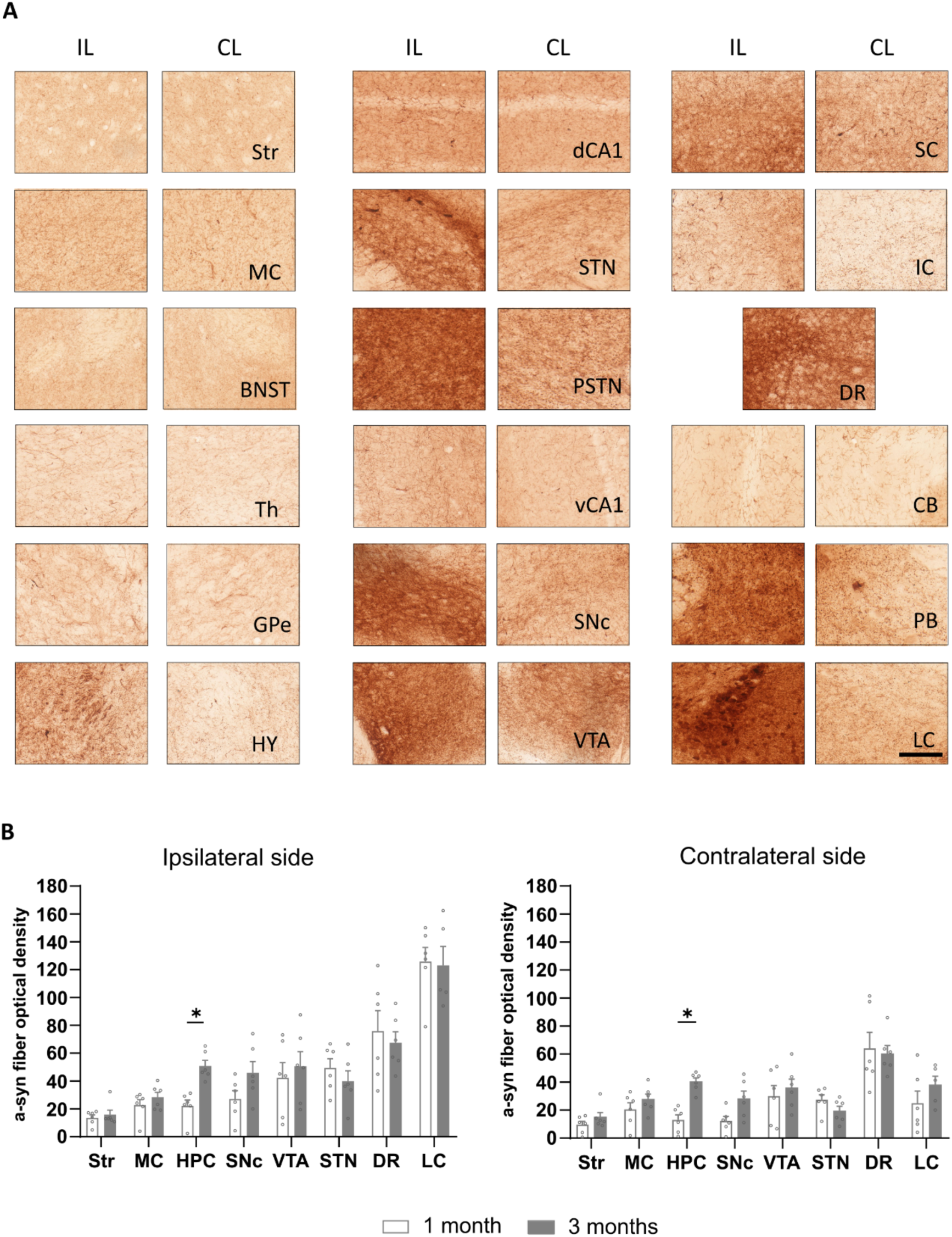
A-syn expression in unilaterally injected brains. (**A**) Representative images of different brain regions expressing a-syn in the ipsilateral and contralateral sides relative to the injection site. Scale bar: 100 µm. (**B**) A-syn fibre density comparing 1 and 3 months. *p < 0.05, Student’s t-test. Str = striatum; MC = Motor cortex; BNST = Bed nucleus of the stria terminalis; Th = thalamus; GPe = external globus pallidus; HY = hypothalamus; dCA1 = dorsal CA1; STN = subthalamic nucleus; PSTN = parasubthalamic nucleus; vCA1 = ventral CA1; SNc = Substantia nigra pars compacta; VTA = ventral tegmental area; SC = superior colliculus; IC = inferior colliculus; DR = dorsal raphe; CB = cerebellum; PB = parabrachial nucleus; LC = Locus coeruleus; HPC = hippocampus. N = 7 animals/group

### A-Syn accumulation preserves LC and SNc neurons while affecting projection areas

To determine whether a-syn overexpression induces cell loss, we performed immunofluorescence staining for TH and quantified noradrenergic neurons in the LC. In a-syn mice, no significant reduction in TH+ neurons was observed at either sex or time point (Fig. 4A). Given the central role of the SNc in PD pathology, we also assessed dopaminergic neuron density in this region using stereological methods. No significant differences were observed between sham and a-syn mice, regardless of sex or time point (Fig. 4B). To investigate whether a-syn expression impacts noradrenergic terminals, we quantified DBH+ fibres in the HPC. Our results showed a significant decrease in DBH+ fibre density in the HPC of male a-syn animals, with a reduction also evident at 1 month in a-syn females (Fig. 4C).

**Figure 4.**
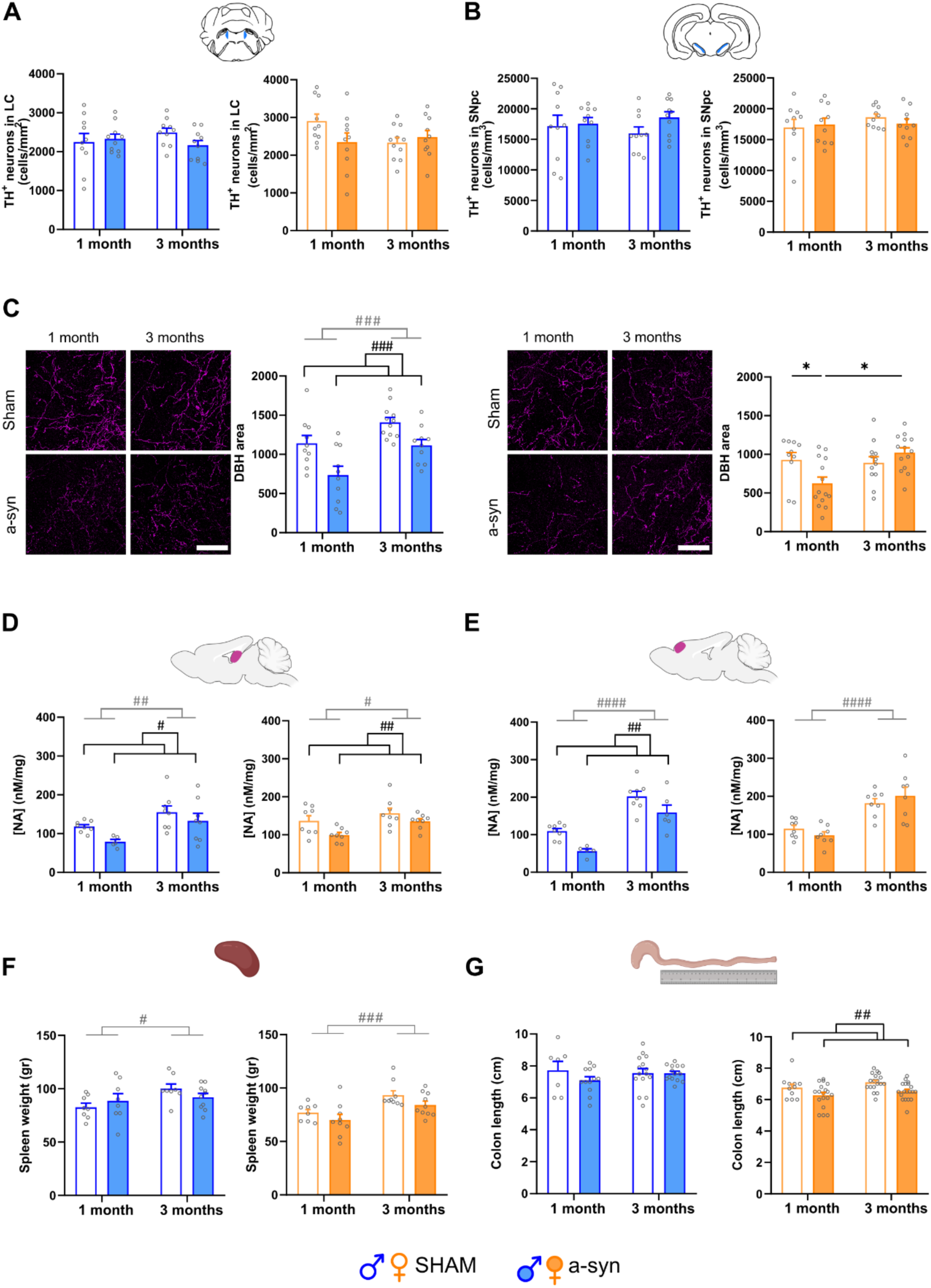
A-syn impact in brain and periphery. (**A**) TH+ neuron quantification in the locus coeruleus shows no changes between sham and a-syn animals. N = 5 animals/group (**B**) Stereological analysis of SNc neurons reveals no degeneration. N = 5 animals/group (**C**) DBH fibre quantification in the hippocampus showed reduction in a-syn mice. Scale bar: 50 µm (40x). N = 5-6 animals/group (**D**) HPLC analysis revealed decreased noradrenaline levels in the hippocampus and (**E**) prefrontal cortex in a-syn mice. N = 4 animals. (**F**) No differences were found in spleen weight between sham and a-syn. N = 8-10 animals/group (**G**) Female a-syn mice exhibited shorter colon length. N = 7-19 animals/group # p < 0.05, ## p < 0.01, Two-way ANOVA; *p < 0.05, Tukey’s post-hoc test.

Next, NA levels were measured via HPLC analysis in the HPC, PFC and general cortical region. In HPC, male a-syn animals displayed a significant reduction in NA levels compared to sham in both sexes (Fig. 4D). In the PFC, NA levels were also reduced in males but not in females (Fig. 4E). This trend was also observed in general cortical area, where a-syn males showed reduced NA concentrations, whereas in females this difference was not detected (Supplementary Fig. 2A). Given that serotonin (5-HT) alterations are also implicated in the prodromal phase of PD and that the LC modulates serotonergic transmission,^50–52^ we further analysed 5-HT levels by HPLC. In the HPC, 5-HT was reduced only in a-syn males but not in a-syn females (Supplementary Fig. 2B). In the PFC, both sexes exhibited a significant reduction in 5-HT levels in the a-syn group (Supplementary Fig. 2C). In contrast, in the cortex, 5-HT levels remained unaffected in both sexes (Supplementary Fig. 2D). These findings indicate that a-syn overexpression disrupts NA and 5-HT homeostasis in LC-projection areas in a region and sex-dependent manner.

To explore potential systemic alterations linked to the a-syn model, we assessed spleen weight and colon length as indicators of systemic inflammation and gut-brain axis dysfunction respectively. Spleen weight remained unchanged in a-syn animals, but an age-related increase was observed at 3 months (Fig. 4F). Regarding colon measurements, while male a-syn mice showed no differences in colon length, female a-syn mice exhibited a significant reduction compared to sham animals (Fig. 4G).

### Sex-dependent functional differences in the LC in the a-syn model

To investigate whether a-syn overexpression induces functional alterations in LC neurons, we conducted patch-clamp recordings (Fig. 5A) at 1 and 3 months post-injection. The results at 1 month revealed sex-dependent alterations in basal firing rates, passive membrane properties and excitability.

**Figure 5.**
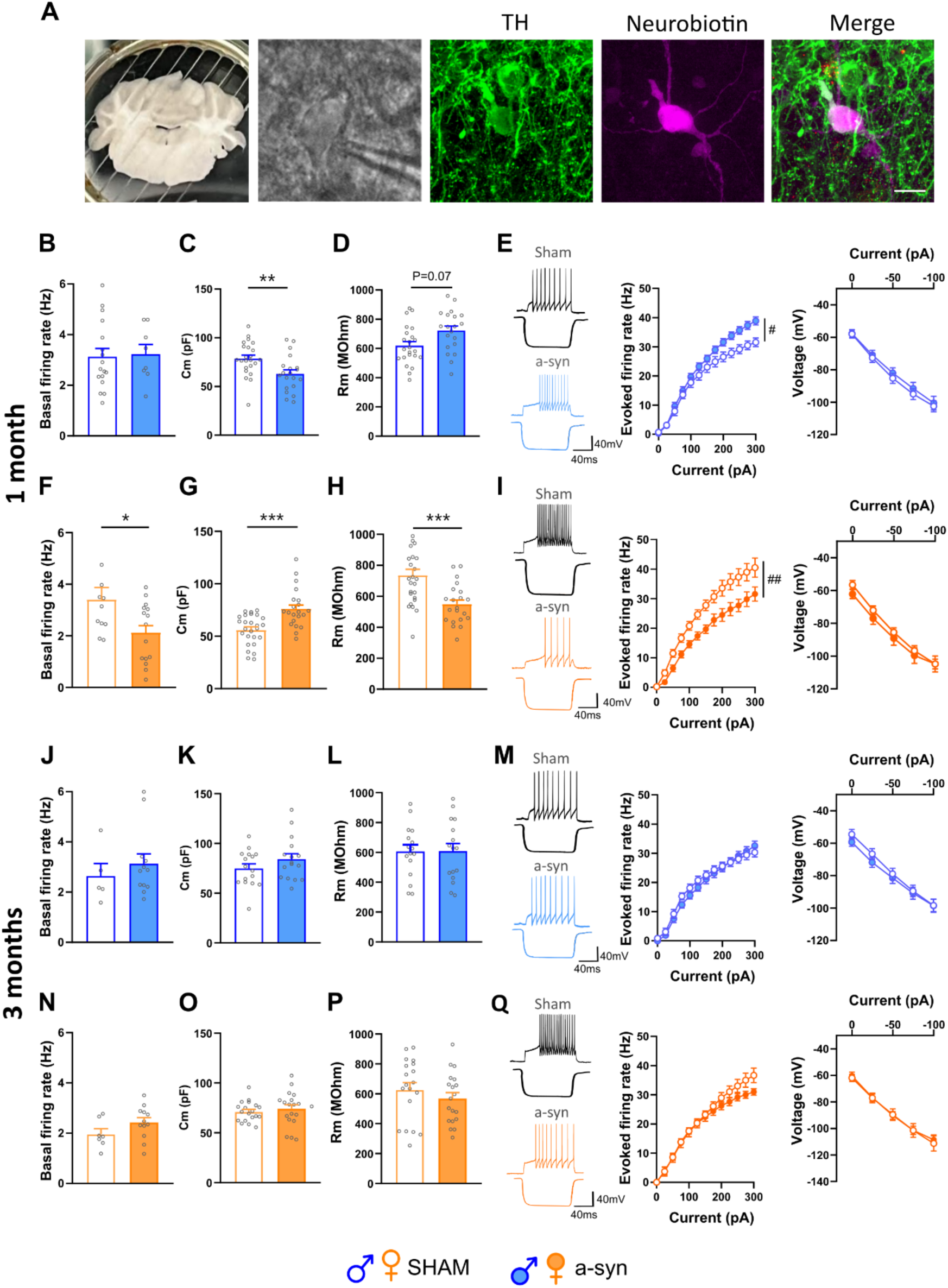
In vitro recordings of locus coeruleus neurons overexpressing a-syn. (**A**) Representative images of locus coeruleus neurons recorded in patch-clamp. (**B–E**) 1 month a-syn males exhibited decreased capacitance and increased excitability. (**F**) A-syn females showed reduced spontaneous firing frequency at 1 month. (**G–I**) 1 month a-syn females displayed increased capacitance and decreased excitability. **(J–Q**) No differences were observed in basal firing rate, intrinsic properties, or excitability between sham and a-syn mice at 3 months. N = 6 animals/group. Student’s T-test *p<0.5, **p<0.01, ***p < 0.001; Two-way ANOVA #p < 0.05, ##p<0.01.

In cell-attached recordings, basal firing rates did not differ in male a-syn mice compared to sham animals (sham 3.13 Hz, a-syn 3.22 Hz) (Fig. 5B). However, female a-syn mice exhibited significantly reduced basal firing rate compared to sham animals (sham 3.41 Hz, a-syn 2.13 Hz) (Fig. 5F). Passive membrane properties also varied among sexes. In males, a-syn expression induced a decrease in membrane capacitance and a trend toward increased membrane resistance (Fig. 5C and Fig. 5D). In contrast, female a-syn mice exhibited the opposite effect, with increased membrane capacitance and decreased membrane resistance (Fig. 5G and Fig. 5H). Voltage-current curves highlighted further sex-dependent differences. While a-syn males showed increased excitability with higher evoked firing rates during depolarizing steps (Fig. 5E), female a-syn mice exhibited reduced evoked firing rates (Fig. 5I). Regarding AP properties, no relevant significant changes were detected (Supplementary Table 3 and Supplementary Table 4). At 3 months post-injection, none of these electrophysiological alterations were observed. Basal firing rates, passive membrane properties, voltage-current relationships, and AP properties were comparable between a-syn and sham animals in both sexes (Fig. 6 J-Q).

**Figure 6.**
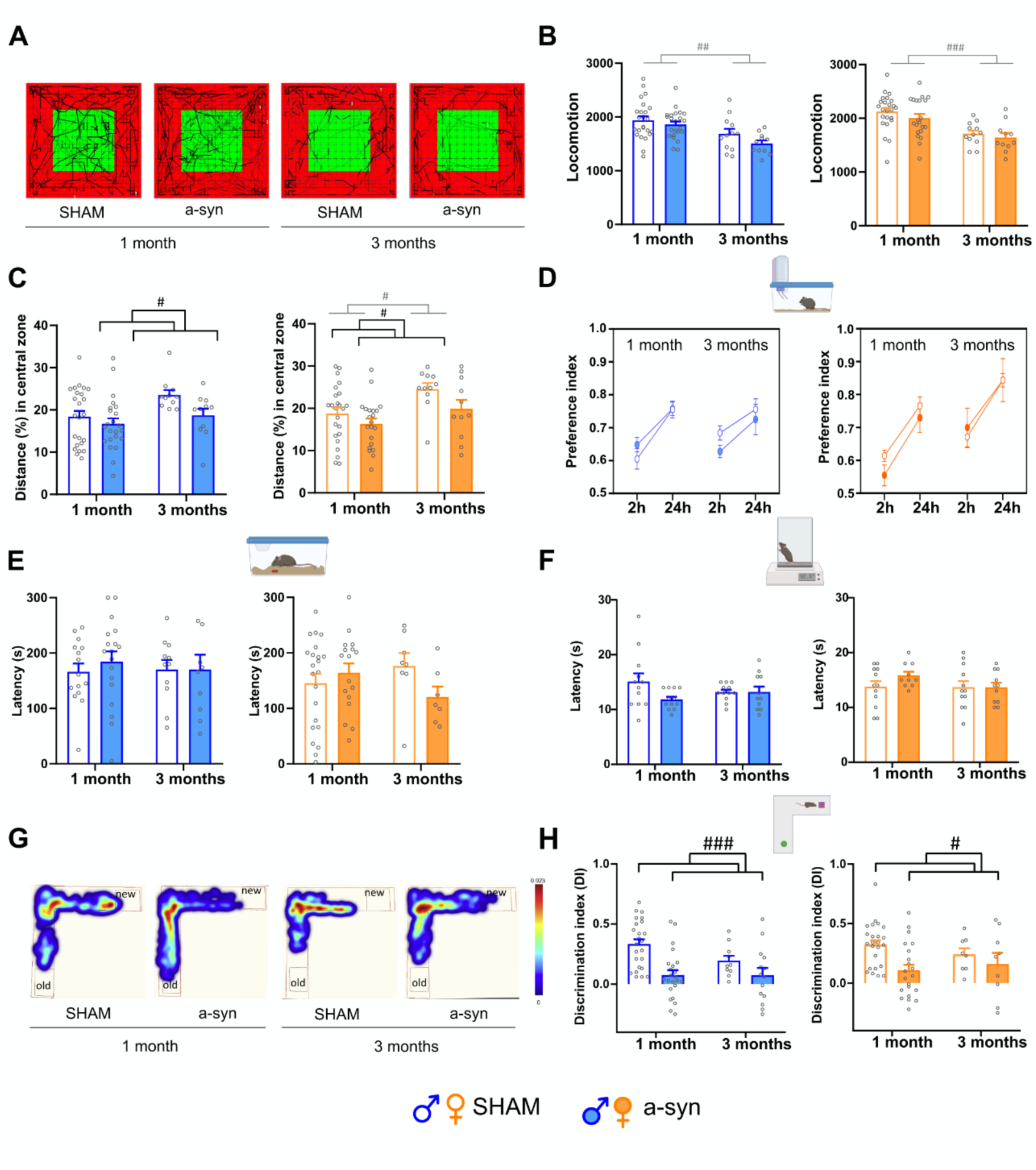
Behavioural evaluation of the a-syn model. (**A**) Representative images of open field performance. (**B**) No motor differences were observed between sham and a-syn mice. (**C**) A-syn overexpression reduced the distance percentage in the centre of the arena, indicating anxiety-like behaviour in both sexes. No differences were found in anhedonia (**D**), olfaction (**E**), or nociception (**F**). (**G**) Heat maps of exploratory behaviour in the novel object recognition test. (**H**) Discrimination index was significantly reduced in a-syn males. N = 10-24 animals/group. # p < 0.05, ### p < 0.001, Two-way ANOVA.

Together, these findings indicate that a-syn expression induces transient, sex-dependent alterations in LC electrophysiology, with functional differences evident at 1 month but not sustained at later stages (Fig. 5 J-Q).

### A-syn Overexpression Induces Cognitive Deficits and Mild Anxiety-Like Behaviour

To determine whether the observed LC functional alterations translate into behavioural outcomes, we next assessed motor, affective, and cognitive functions in a-syn mice.

Our results suggested preserved motor function with no significant differences in locomotion (Fig. 6A and Fig. 6B), general activity or stereotypic movements (Supplementary Fig. 3A-D). This finding aligns with the absence of cell loss in the SNc and is consistent with the premotor phase of PD, where motor symptoms are absent.

Given the relevance of NA in neuropsychiatric symptoms appearance in PD,^14,30,31^ we evaluated anxiety-like behaviour and anhedonia. In the OF test, a-syn mice of both sexes travelled significantly less in the centre of the arena (Fig. 6C), indicating increased anxiety. This was further supported by the dark and light box test were female a-syn mice exhibited shorter latencies to enter the dark zone whereas no significant differences were observed in males (Supplementary Fig. 3E-H). However, no differences were observed in the elevated plus maze (Supplementary Fig. 3I-L). Sucrose preference, did not differ between a-syn and sham animals, indicating no clear signs of anhedonia (Fig. 6D). Olfactory function was evaluated using the buried food task, revealing no significant differences between a-syn and sham animals (Fig. 6E). Similarly, nociceptive responses in the hot plate test showed no overall differences (Fig. 6F). Interestingly, cognitive deficits were evident in the a-syn mice, as shown in the NOR task. A-syn animals displayed robust lower discrimination indices, regardless of sex (Fig. 6G and Fig. 6H).

Overall, behavioural evaluation revealed anxiety and cognitive deficits in male and female a-syn mice. Importantly, these behavioural deficits were persistent and remained 3 months after AAV injection.

## Discussion

The contribution of noradrenergic dysfunction to PD pathogenesis remains unclear, largely due to the lack of models that selectively target the LC. Here, we described a novel mouse model that overexpresses a-syn in LC neurons, reproducing key prodromal features including neuronal a-syn accumulation, axonal pathology, monoaminergic disruption, sex-dependent electrophysiological alterations, cognitive deficits, and anxiety, without overt neuronal loss. This model uniquely isolates LC-driven pathology, providing a platform to dissect circuit-specific mechanisms and translational interventions.

Using the PRSx8 promoter, we achieved selective a-syn expression in TH+ LC neurons with high efficiency.^53^ Phosphorylated a-syn, a hallmark of early aggregation and Lewy body formation,^49,54^ was detected in somas as early as one month after AAV-PRSx8-hasynWT injection, while monomeric a-syn propagated extensively along LC axons. Consistent with previous studies using similar a-syn carrying-viral vectors,^55–59^ soluble a-syn species propagated to projection areas including the DR, HPC, PFC and SNc largely colocalizing with NET+ fibres, suggesting a-syn axonal transport.^60–62^ The presence of a-syn in some regions from the contralateral hemisphere, as the HPC, aligns with the bilateral innervation of LC projections to certain areas^55,63,64^ and with the evidence that commissural pathways may facilitate the interhemispheric communication.^2,65^ Supporting this, animal models have provided compelling evidence that a-syn can propagate to anatomically connected brain regions.^66^ Some authors have also suggested that once a-syn reaches projection areas, it can spread to other interconnected nuclei through prion-like propagation, as described by Braak et al.^3^ In our model, despite robust a-syn accumulation, LC and SNc neuronal loss was absent, consistent with the clinical observations that LC neurons can survive years despite a-syn pathology.^67^ Reduced DBH+ fibres and decreased NA in LC projection areas, indicate a “dying-back” process where axons degenerate prior to soma loss.^14,68^ This pattern mirrors observations in SNc-targeted a-syn models, ^69–71^ and in toxic-based models of noradrenergic damage using DSP-4, neuromelanin, or 6-OHDA or DBH-hSNCA mice,^42,44,72,73^ further supporting selective axonopathy in alpha-synucleinopathies.^74^ In addition to NA content loss, serotonergic transmission was also affected in our a-syn model, as happens in patients^75^ and some mouse models of the disease.^76^ The fact that the LC influences DR activity has been well described in the literature. The LC extensively innervates the DR^77,78^ modulates its excitability,^51^ and compromises 5-HT release.^50^ This interplay highlights the LC as a hub integrating monoaminergic balance across cortical and limbic targets.

In addition to neuroanatomical changes, recent findings increasingly support the bidirectional communication along the gut-brain axis.^2^ In our study, colon shortening, observed in female a-syn mice, suggests early enteric involvement, consistent with gut inflammation or dysbiosis observed in PD patients.^79^ Absence of abnormalities in spleen weight suggests minimal systemic inflammation.^80^ These results align with “brain-first” progression, as accumulating evidence, including our own, supports a-syn spread from central to enteric regions via vagal pathways.^66,81^ Notably, the female-specific phenotype mirrors clinical reports of greater gastrointestinal symptoms in women,^82,83^ supporting a potential sex-dependent vulnerability of the enteric nervous system in PD.

Some authors suggest that, in the early stages of PD and Alzheimer’s, LC neurons become hyperexcitable due to a-syn overexpression, exhibiting increased firing rates that may precede subsequent hypofunction and degeneration.^67^ In our experiments, however, we observed a reduction in firing frequency exclusively in females, with no changes in males. Previous studies overexpressing wild-type human a-syn in the LC of male mice reported no alterations,^84^ while other LC-targeted models have shown variable outcomes. For instance, 6-OHDA injections and neuromelanin overexpression increased basal firing and pattern of LC neurons,^85,86^ whereas DSP-4 intraperitoneal administrations did not affect tonic activity.^42,87^ Classical PD models targeting the medial forebrain bundle or the SNc, using 6-OHDA,^88,89^ rotenone,^90^ MPTP,^91^ and preformed fibrils^92^, typically report compensatory reductions in spontaneous LC and SNc neuron activity. Importantly, most of these studies were conducted in males or pooled sexes,^42,86^ potentially masking sex-specific responses. The mechanisms underlying the reduced firing activity in a-syn females remain unclear and warrant further investigation. Our AP analyses revealed no changes in afterhyperpolarization or other electrophysiological parameters, suggesting that major ion channel dysfunction is unlikely. Nonetheless, subtle changes in L- and T-type Ca²⁺ channels ^93^ and enhanced GABAergic inhibition, cannot be ruled out.^94,95^ Beyond firing frequency, we also found sex-dependent differences in intrinsic membrane properties. A-syn females displayed higher membrane capacitance, whereas a-syn males showed lower capacitance relative to their respective controls. These opposing effects were accompanied by distinct excitability profiles where neurons from males exhibited hyperexcitability and in females were hypoexcitable. The increased excitability observed in a-syn males may result from smaller neuronal size, as indicated by their reduced capacitance.^96,97^ This may be further exacerbated by the more pronounced monoamine depletion found in males, which likely reflects a greater compromise of LC projections. Although SK-channel dysfunction has been implicated in a-syn induced hyperexcitability,^84^ our data do not support this mechanism, as afterhyperpolarization amplitude was unchanged. Alterations in receptor signalling might also contribute; for example, CB1 receptor deletion in the LC selectively increases excitability in male neurons while leaving females unaffected.^98^ In contrast, the hypoexcitability observed in females may relate to increased membrane capacitance, supporting the idea that a-syn overexpression drives distinct morphological and functional adaptations in each sex.

Our electrophysiological findings highlight the importance of considering sex as a biological variable when studying LC physiology, both under normal and pathological conditions. Sex dimorphism is a well-established feature of the LC in control mice, with females exhibiting greater neuronal number, enhanced dendritic complexity, and distinct electrophysiological and molecular profiles.^99–104^ Under pathological conditions, as shown here, both sexes can undergo divergent alterations. For instance, in a neuropathic pain model using chronic constriction injury, LC neurons from female mice displayed hypoexcitability, whereas those from males were hyperexcitable,^105^ mirroring the changes induced by a-syn in our study. Interestingly, the functional alterations observed in our model appeared to be transient, as most differences had resolved by 3 months. This apparent recovery may reflect compensatory neuronal plasticity, similar to that described in other PD models.^92^

Given the involvement of the LC in PD symptomatology, we next assessed whether its selective a-syn overexpression could lead to behavioural alterations, following the anatomical and functional analyses. Motor impairment, a hallmark of PD, results from the degeneration of dopaminergic neurons in the SNc, which can be exacerbated by LC dysfunction.^106–109^ In our model, motor function, assessed in the OF, was preserved, consistent with the anatomical findings showing that dopaminergic neuron number in the SNc remained unaffected despite elevated a-syn levels. This observation aligns with previous studies employing LC-targeted a-syn expression or DSP-4 lesions, where also locomotor activity was unaltered.^44,86^ Other non-motor behaviours were similarly unchanged, as anhedonic behaviour, olfaction or pain. Although reductions in 5-HT and NA levels were observed, a neurochemical feature often linked to major depressive disorder,^110^ these changes did not translate into behavioural deficits. Methodological factors, such the sucrose concentration^111^ or the absence of deprivation protocols, may have contributed to this outcome. Likewise, no olfactory impairment was detected, possibly influenced by the lack of food deprivation^112,113^ or by the limited time window for a-syn propagation from the LC to the olfactory bulbs.^81,114^ Finally, thermal nociception thresholds remained unchanged, in agreement with prior evidence suggesting that heat sensitivity is often spared in PD,^115^ and that noradrenergic dysfunction may selectively influence other pain modalities.^116–118^

In contrast to the preserved motor and non-motor functions described above, anxiety and cognition were notably affected at all evaluated time points. Anxiety is a common non-motor symptom of PD, affecting up to 60% of patients and often manifesting as generalized anxiety, panic attacks, or social phobia.^119^ In our model, a-syn animals displayed mild but consistent anxiety-like behaviour, reflected by reduced exploration of the centre in the OF. This phenotype may be linked to the decreased NA levels observed in the PFC of a-syn animals, a region critically involved in anxiety regulation. Supporting this interpretation, inhibition of the LC-PFC pathway has been shown to increase anxiety-like behaviour in stressed rodents.^120^ These observations align with anxiety phenotypes reported in neuromelanin transgenic models^121^ and other a-syn models, including DBH-hSNCA mice,^44^ raphe-targeted overexpression models,^56^ and transgenic lines.^76^ In addition to anxiety like behaviour, cognition was also impaired in our animals. On the other hand, cognitive impairment affects up to 83% of PD patients^122^ and, although traditionally linked to cholinergic and dopaminergic dysfunction, growing evidence points to noradrenergic degeneration, particularly LC dysfunction, as a major contributor.^5,16,32,34,123–125^ In our a-syn model, both male and female animals exhibited significant cognitive deficits in the NOR task. The 24-hour retention interval employed strongly engages hippocampal-dependent long-term memory,^126–131^ suggesting that the observed deficits reflect hippocampal rather than perirhinal dysfunction. Consistent with this, NA levels were reduced in both the HPC and PFC, areas densely innervated by the LC,^132,133^ which co-releases NA and DA^134,135^ and plays a central role in novelty detection, attention, and cognitive flexibility.^136–138^ Moreover, a-syn expression was evident in noradrenergic hippocampal projections, suggesting direct involvement of LC-derived fibres in local circuit alterations. Given that the LC bilaterally innervates the HPC, even restricted unilateral a-syn overexpression is likely to compromise hippocampal integrity, a notion supported by comparable cognitive impairments described in other a-syn models.^139–141^ Collectively, these data indicate that selective a-syn overexpression in LC neurons is sufficient to trigger specific anxiety-like and cognitive deficits, while sparing motor performance. This pattern mirrors the prodromal phase of PD, in which early noradrenergic dysfunction contributes to emotional and cognitive disturbances preceding overt motor decline.

In conclusion, selective overexpression of a-syn in noradrenergic LC neurons recapitulates key features of prodromal PD, including somatic a-syn accumulation, widespread axonal distribution, and monoaminergic dysfunction without overt neuronal loss. Behaviourally, the model reproduces key early non-motor features as mild anxiety-like behaviour and marked cognitive impairments in both sexes, in the absence of motor deficits, closely mirroring the clinical progression of early PD. These findings highlight the LC as a pivotal node linking noradrenergic imbalance to emotional and cognitive disturbances, establishing a robust experimental platform to investigate the circuit-level consequences of early a-syn pathology. The presence of sex-specific and transient alterations in neuronal excitability underscores the importance of considering sex as a biological variable when developing LC-targeted therapies for PD. From a translational perspective, the demonstration of axonal a-syn propagation and persistent NA depletion supports the LC as a strategic target for early intervention aimed at preserving circuit integrity before irreversible neurodegeneration occurs. Furthermore, the peripheral alterations detected, including early enteric involvement in females, emphasize the need for integrated central–peripheral biomarker approaches to improve early PD diagnosis and to guide the development of disease modifying therapies.

## Acknowledgements

The authors thank for technical and human support provided by SGIker (UPV/EHU/ ERDF, EU) Core Facility for Analytical and High-Resolution Microscopy in Biomedicine, and Bizkaia General Animal Facility. Africa Vales for the construction and production of the AAV vectors.

## Supplementary material

Supplementary material is available at *Brain* online.

## Supplementary Figures

**Supplementary Fig. 1.**
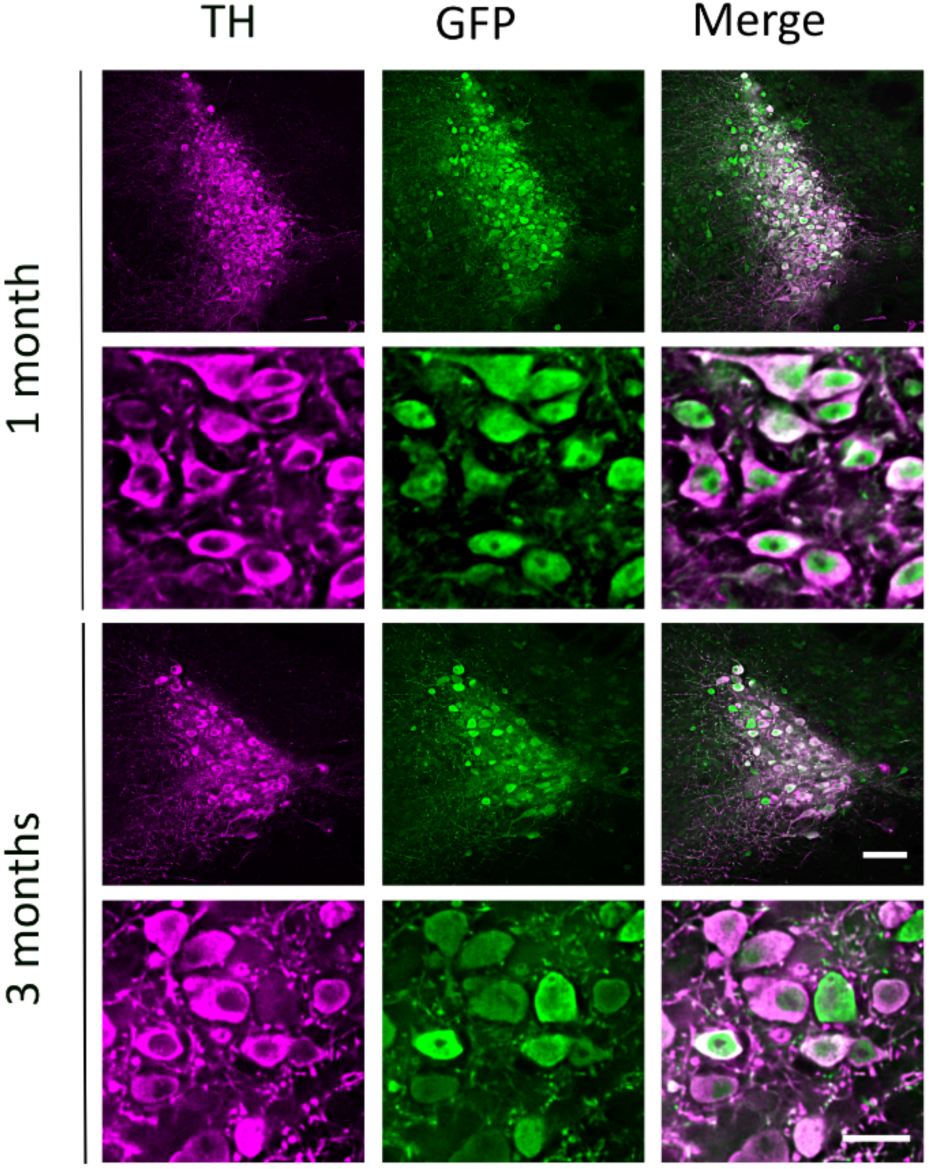
TH-GFP immunofluorescence in the Locus Coeruleus. GFP-stained neurons colocalize with TH+ neurons of the locus coeruleus. Images were captured at 20x magnification (scale bar 100 µm) and 63x magnification (scale bar 25 µm).

**Supplementary Fig. 2.**
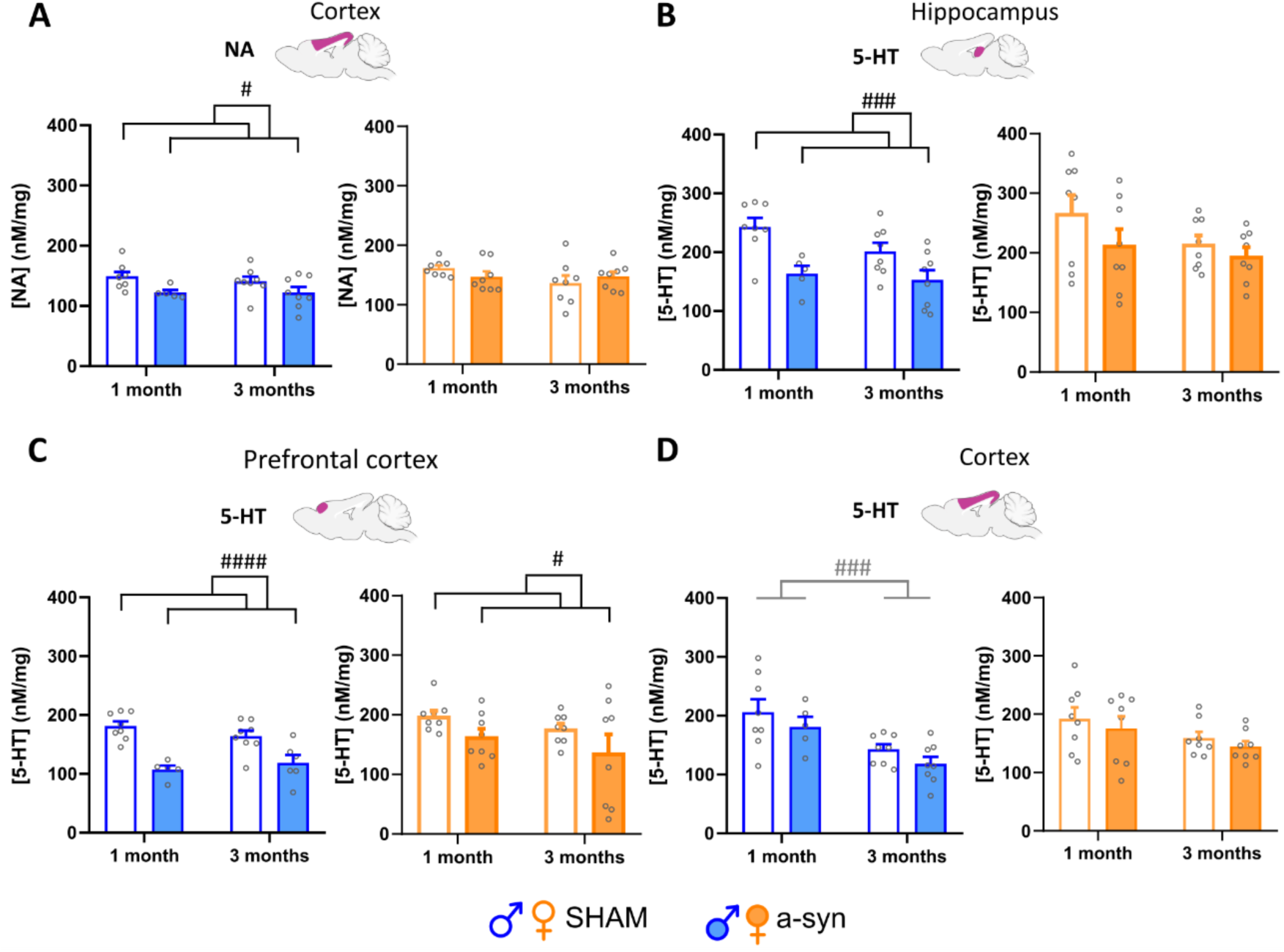
HPLC measurements of noradrenaline and serotonin. (**A**) Noradrenaline levels were reduced in the cortex of male a-syn mice, with no significant changes observed in females. (**B**) In the hippocampus, serotonin levels were reduced in a-syn males but not in females. (**C**) Serotonin levels in the prefrontal cortex were decreased in both sexes in the a-syn group. (**D**) No differences in cortical serotonin levels were found between sham and a-syn mice.

**Supplementary Fig. 3.**
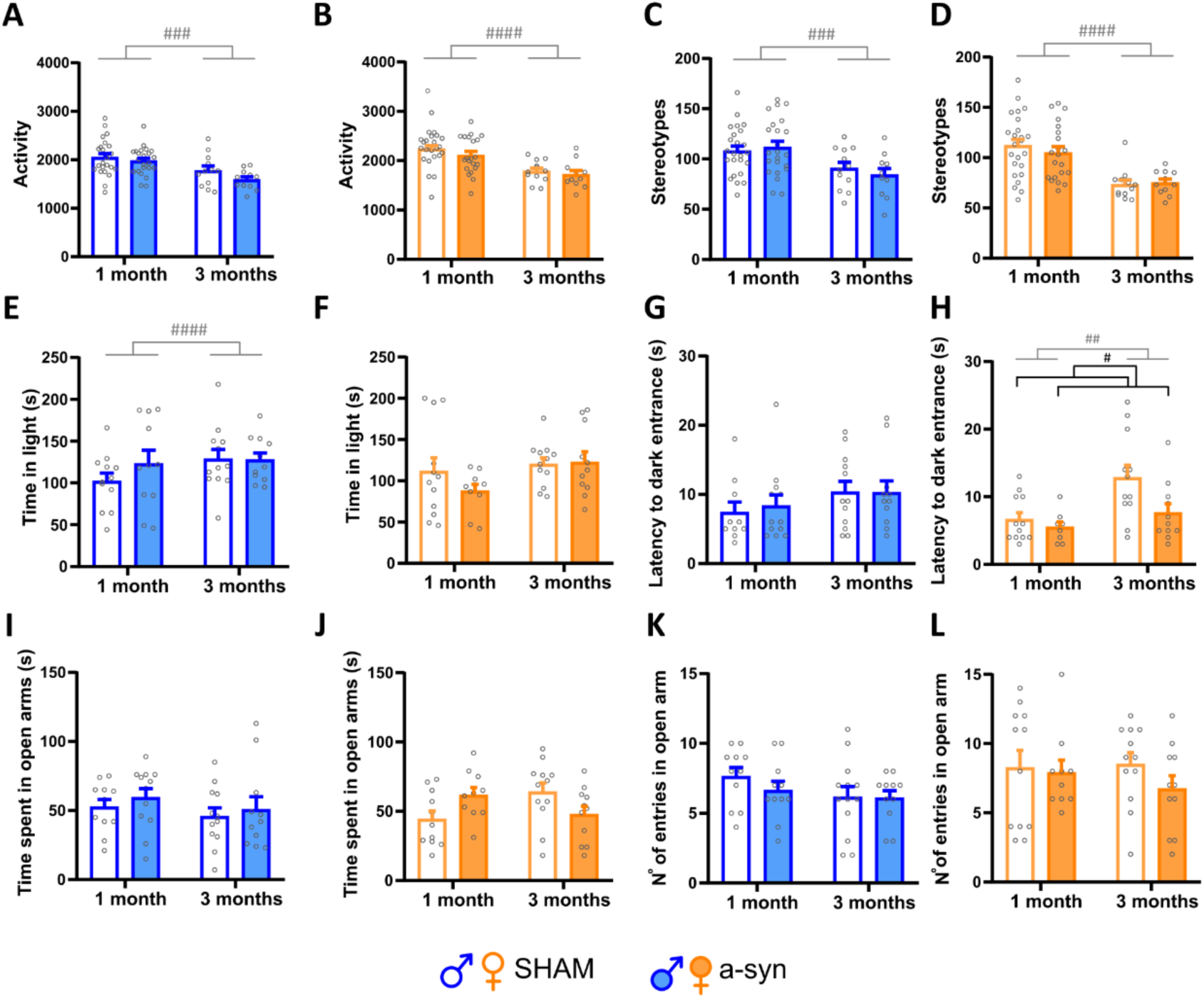
Behavioural analysis of the a-syn model. (**A-D**) Motor activity and stereotypy analysis in the Open Field test showed no differences among the groups. (**E-H**) Dark and Light Box test showed that a-syn mice took less time to enter to the dark compartment. (**I-L**) No differences were observed between groups in the Elevated Plus Maze test.

## Supplementary information

### Animals

131 male and 117 female C57BL/6J mice (Envigo, Spain), 8 weeks old at the beginning of the experiment, were used in this study. Animals were housed in groups of six under standard laboratory conditions (22 ± 1°C, 55 ± 5% humidity, and a 12:12 h light/dark cycle) with *ad libitum* access to food and water. Animal weight was monitored weekly. All procedures followed ARRIVE guidelines and were conducted in compliance with EU Directive 2010/63/EU and Spanish regulations (RD 53/2013) for laboratory animal care. Protocols were approved by the Local Committee for Animal.

### Viral vector and stereotaxic surgery

Animals were injected bilaterally or unilaterally in the locus coeruleus (LC) with different adeno-associated viral vectors encoding the hSNCA gene. A viral construct with a specific promoter for DBH neurons was used, using the following viral vectors: AAV9-MCS-PRSx8-GFP-WPRE, AAV9-MCS-PRSx8-WPRE, and AAV9-MCS-PRSx8-hasynWT-WPRE. A total volume of 0.5 µL of viral vector (1.07 × 10¹³ gcp/mL) was injected into each LC. All viral constructs were produced at the CIMA (Centro de Investigación Médica Aplicada, Universidad de Navarra, Spain) facilities in collaboration with Dr. Gloria González Aseguinolaza.

Animals were anesthetized with isoflurane (4% for induction and 1.5-2% for maintenance) and oxygen 0.8-1%. They were placed in the stereotaxic frame (David Kopf® Instruments, Tujunga, California, EEUU, model 957). A heating pad was placed for maintaining the body temperature at 33 °C and lubrithal® was used for eye lubrication. Meloxicam (2 mg/kg) was injected subcutaneously to reduce pain and inflammation. The scalp was incised, the skull exposed. A hole of 0.5-1 mm in diameter was drilled at the following coordinates relative to the skull: from bregma, anteroposterior = −5.4 mm, mediolateral = ±0.9 mm, dorsoventral = −3.5 mm (Paxinos & Franklin, 2004). The viral vector was infused into each hemisphere at a rate of 0.2 μl/min. Injections were performed using pre-pulled pipettes (Blaubrand IntraMARK 708707, Germany) attached to a syringe to exert pressure. The needle was left in place for 5 minutes post-injection to ensure proper viral diffusion.

### Immunohistochemical procedures

#### Tissue processing

Animals were deeply anesthetized with ketamine (100 mg/mL) and xylazine 2% (at doses of 100 mg/kg and 10 mg/kg, respectively) and then transcardially perfused with 4% paraformaldehyde. Brains were fixed with 4% paraformaldehyde during 24h and dehydrated with 30% sucrose and 0.1% sodium azide in phosphate buffer 0.1M solution until the brains were sunk and prepared for being cut. Brain tissue was sectioned on a freezing microtome (Microm 440E, Germany) at 40 μm and stored in cryoprotectant solution (0.157% NaH2PO4, 1.36% Na2HPO4, 30% ethylene glycol+ 26% glycerol) at −20°C until further processing.

When immunohistochemical verifications were performed on slices previously recorded using ex vivo electrophysiology, the 220-micrometer thick slices were fixed for 24 hours in 4% paraformaldehyde. Following fixation, the slices were washed with 0.1M PBS three times for 5 minutes each before being placed in a cryoprotectant solution.

#### Immunohistochemistry with 3’3 Diaminobencidine (DAB)

DAB immunoassays were performed for revealing TH in the SNc slices, as well as for a-syn detection thorough the brain axis.

Free-floating sections were treated to block endogenous peroxidase activity with 0.9% H₂O₂ in PBS for 20 minutes, followed by washing with 0.1 M PBS. The sections were then incubated in a blocking solution (0.1 M PBS + 5% NGS + 5% Triton X-100) for 1 hour. Subsequently, the samples were incubated overnight with rabbit anti-TH (Supplementary Table 1) at room temperature in the same blocking solution. The sections were washed with 0.1 M PBS buffer and incubated for 2 hours shaking with the secondary antibody (Supplementary Table 1). Antigen signal was amplified using VectaStain® Elite ABC-HRP Kit 1:100 (PK-6100, Vector laboratories) + 0.1 M PBS + 0.05% Triton X-100 during 1 hour at room temperature. The staining was revealed using 2% 3’3-diaminobenzine (DAB) in PBS 0,1M + triton X-100 0,5% solution and 3% H_2_O_2_. Once the immunohistochemical reaction was performed, the slices were placed on Superfrost slides. Finally, the samples were dehydrated in xylol at least 2 hours and covered with DPX mounting medium.

#### Immunofluorescence staining

Initially, the sections were rinsed twice with 0.1 M PBS for 5 minutes each to remove cryoprotective solution. They were then preincubated for 2 hours in a solution containing 5% NGS and 0.3% Triton X-100 in PBS. Next, the sections were incubated overnight at 4°C with the appropriate primary antibodies (Supplementary Table 1), diluted in a solution of 5% NGS, 0.3% Triton X-100, and PBS. The following day, they were incubated with secondary antibodies (Supplementary Table 1) for 2 hours in a solution of 0.3% Triton X-100, 1% NGS, and 0.1 M PBS. Finally, the sections were mounted onto Superfrost slides and covered using Vectashield mounting medium (Vector Labs, Peterborough, UK). For DBH immunofluorescence, an antigen retrieval procedure was used. Briefly, slices were incubated in 2N HCl for 15 minutes at room temperature with shaking. They were then washed three times with 0.1M PBS, and the standard immunofluorescence protocol was followed.

#### Quantification and analysis

Fluorescent images were captured using a Zeiss Apotome microscope with structured illumination or a Zeiss LSM880 Fast Airyscan super-resolution confocal microscope for qualitative analysis. The images were then edited using ImageJ software (version 1.8.0, NIH).

### TH+ and a-syn neuron semiquantification and colocalization in locus coeruleus

Noradrenergic neurons of the LC were quantified using TH immunofluorescence, while a-syn staining was employed to quantify colocalization with TH+ neurons.

LC neurons were quantified from a total of 5 mice per group. 3 slices from each mouse were selected, one every third section in the rostromedial extent from bregma −5.34 to - 5.52 mm. Double immunofluorescence staining for TH and a-syn was performed as previously described. The images were captured using a Nikon Ti-U conventional fluorescence microscope equipped with a Nikon Ds-Qi2 camera (Nikon, Japan) from de SGIker facilities of the UPV/EHU. Images were acquired at 20x magnification (CFI Plan Apo Lambda 20x/0.75 WD 1.0 mm), with a Z-stack acquired at a step size of 0.9 µm. To enhance image sharpness and clarity, Richardson-Lucy deconvolution was applied. ImageJ software was used to adjust brightness and contrast, and to create maximum intensity projection (MIP) images. TH expression and TH-a-syn colocalization were manually counted by an experimenter who was blind to the experimental conditions.

Neuron colocalization was expressed as efficiency, defined as the ratio of neurons positive for both a-syn and TH to the total number of TH+ neurons (N (a-syn/TH)/ N (TH)). The number of neurons was expressed as density, quantified as TH+ neurons per mm².

#### Substantia nigra pars compacta neuron stereological quantification

Five animals per group were chosen for this experiment and each hemisphere was analysed separately, 7 slices between bregma −2.92 mm and −3.16 mm were used for the analysis, letting 80 mm distance between slices, according to Paxinos and Watson atlas. Boundaries of the area of the SNc were traced as a ROI (region of interest) with the 4x objective. The sections were quantified using a counting frame size 50x50μm and a distance of 60μm x 60μm between dissectors.

The Mercator image analysis system (Explora-Nova, La Rochelle, France) was used along with a digital camera connected to an Olympus BX51 microscope, which features a three-axis motorized stage. The optical fractionator method involves estimating the total number of cells based on the number of cells sampled within a set of equidistant virtual counting spaces in the X, Y, and Z directions, without any bias. To calculate the total number of cells, the following formula was employed: N = ΣQ x (1/ssf) x (1/asf) x (1/hsf), where Q represents the actual number of cells counted in a sample, and N is the total cell estimation. In this study, the section sampling fraction (ssf) was 1/3. Volume estimations were obtained using Cavalierís method.

#### Optical density of a-syn expression in brain areas

To analyze a-syn overexpression in specific brain regions, unilateral injections were performed on male mice (n=7 for each condition). A-syn overexpression was assessed using DAB immunostaining. These sections were captured using high resolution slide scanner Pannoramic MIDI II (3DHistech, Hungary) from the Achucarro Basque Center for Neuroscience. To determine the a-syn overexpression in different brain areas, optical density analysis was conducted. In each section, the region of interest (ROI) was delineated, and the average optical density of the hemisphere was compared with an area without a-syn in the same slice, which served as the background reference. The ROIs analyzed with this method included the striatum, motor cortex, hippocampus (HPC), SNc, ventral tegmental area, subthalamic nucleus, dorsal raphe, and LC. The images were analyzed using FIJI software ^7^.

### Brain Tissue NA and 5-HT Analysis

Mice were euthanized by cervical dislocation following ketamine-xylazine anaesthesia (100 mg/mL / 2%). Brains were rapidly dissected, isolating the HPC, the general cortex and the prefrontal cortex (PFC) which were frozen until further processing. The caudal brain was used for a-syn-DAB immunohistochemistry to confirm LC overexpression. Tissue was fixed (4% paraformaldehyde, 24 h), transferred to a 30% sucrose solution with 0.1% sodium azide, and sectioned at 40 µm. Samples were weighed, treated with 1N perchloric acid and 100 μM EDTA, then homogenized (Ultra-Turrax® T-10) and centrifuged (15 min, 4°C, 20,817 × g). The supernatant was filtered (Costar Spin_XTM 0.22 μm, Sigma-Aldrich) by a second centrifugation (5 min, 4°C, 1,000 × g). Monoamine concentrations were measured using HPLC with electrochemical detection (VT-03, Antec Scientific) at 0.3V. Separation was achieved with a C18 column (ALF-215, 2.1 × 150 mm, Antec Scientific) and a mobile phase containing 50 mM phosphoric acid, 0.1 mM EDTA, 8 mM NaCl, 500 mg/L sodium octyl sulfate, and 12% methanol (pH 6.0, 0.2 ml/min flow rate). Chromatographic data were analysed using Chemstation plus software, and monoamine concentrations were determined via linear regression of standard curves (0.5–400 nM).

### In vitro electrophysiology

Mice were anesthetized with 5% isoflurane, then decapitated, and their brains were rapidly removed and kept in cold. Coronal sections (220 μm) containing the LC were cut using a vibratome (Microm HM 650V) in cold, sucrose enriched artificial cerebrospinal fluid (ACSF) (2.5 KCl, 1.25 NaH₂PO₄, 2.5 KCl, 0.5 CaCl₂, 10 MgSO₄, 10 glucose, 230 sucrose). Slices containing the LC were transferred to warmed (35°C) ACSF at least 30 minutes before being moved to the recording chamber. The ACSF had the following composition (in mM): 126 NaCl, 1.25 NaH₂PO₄, 2.5 KCl, 2 CaCl₂, 2 MgSO₄, 10 glucose, 26 NaHCO₃, 1 sodium pyruvate, and 4.9 L-glutathione reduced. The solution was equilibrated with 95% O₂ and 5% CO₂, maintaining a pH of 7.3-7.4.

Slices were transferred into the recording chamber and perfused continuously with oxygenated modified ACSF (126 NaCl mM, 1.25 NaH₂PO₄ mM, 3 KCl, 1.6 CaCl₂ mM, 1.5 MgSO₄ mM, 10 glucose mM, 26 NaHCO3 mM) heated to 32-34°C (TC-324B Single Channel Automatic Heater Controller, Warner Instruments, USA). LC neurons were identified, using an upright microscope with infrared optics (Eclipse E600FN, Nikon) and a 60X water-immersion objective (60X/1.00 W, Nikon).

For the recording, borosilicate glass capillaries (GC150F-10, Harvard Apparatus) were pulled using a micropipette puller (PC-10, Narishige) to achieve an impedance of 3–6 MΩ. The pipettes were filled with an internal solution containing (in mM): 130 K-Gluconate, 10 HEPES, 5 NaCl, 1 MgCl₂, 1 Mg-ATP, 0.5 Na-GTP, and 10 phosphocreatine disodium salt hydrate (pH 7.4, 280 mOsm). Neurobiotin (SP-1120; Vector Labs), previously stored at 2% in 0.5 M NaCl, was added to the internal solution at a concentration of 1% on the day of the experiment.

Recordings were obtained using an Axopatch 200B amplifier and Digidata 1322A digitizer (Molecular Devices, Sunnyvale, CA, USA) controlled by Clampex 10.3. The image was visualized in a TV monitor

IRK channel activity was recorded in voltage-clamp mode injecting negative pulses The current-voltage curves exhibited characteristic inward rectification currents of LC neurons in both sham and experimental groups.

### Behavioural assessment

#### Open field test

Spontaneous locomotor activity was measured in an open field arena.^1^ The detection of activity was automated using custom-designed software called Actitrack (Panlab, Spain). This device consisted of a square arena (44 x 44 x 35 cm) constructed with Plexiglas and equipped with parallel frames fitted with infrared beams for precise detection of animal movements. The software allowed the arena floor to be divided into two squares to distinguish central and peripheral areas. This configuration was used to detect anxiety-like symptoms in the animals, as anxious individuals tend to spend more time in the periphery of the arena.

For the test, each animal was placed in the centre of the arena, and its activity was recorded for a period of 10 minutes.

#### Elevated plus maze

The elevated plus maze test was employed to evaluate anxiety-like behaviour in mice.^2^ This test exploits the aversion of mice to open spaces and their preference for enclosed areas, providing a measure of their anxiety levels. The apparatus consisted of two open arms (66 × 5.5 cm) and two closed arms (66 × 5.5 cm) arranged in a cross shape on a platform elevated 49 cm above the ground. Each mouse was placed in the centre of the maze facing an open arm and allowed to explore the maze freely for 5 minutes.

#### Dark and light box

To assess anxiety-like behaviour using a more aversive test, the dark and light box test was conducted.^3^ The box (Panlab, Spain) consists of an arena with two separate zones (26 x 26 cm). One side maintains a dim red light and a black floor, while the other side has a brighter light with a white floor. Mice naturally prefer the dark zone and will only venture into the light area when they want to explore. Increased time spent in the light area indicates lower anxiety levels. Each mouse was placed in the centre of the illuminated area facing away from the dark zone and allowed to explore for 5 minutes.

#### Sucrose preference test

To assess anhedonia in mice, the sucrose consumption test was employed, which is a non-invasive and non-stressful method for evaluating depressive-like behaviour in animals. This test measures the animal’s preference for a sucrose solution over regular water, with a reduction in sucrose consumption indicating anhedonia, a core symptom of depression.

Mice were first habituated to drink from two bottles to avoid any side bias before conducting the test. During the test phase, one bottle was filled with water, and the other with a 3% sucrose solution. The placement of these bottles was randomized to prevent positional bias. The mice’s consumption of each liquid was measured by weighing the bottles at 2 and 24 hours.

#### Buried food task

Olfactory impairment of the animals was evaluated using the buried food task.^4^ No food depravation was implemented for this test to avoid interferences with other behavioural tests. Mice were individually housed and familiarized with the scented pellet (Versele-Laga’s Complete Crock®, cheese or apple flavours) by placing the pellet in their cages for 72 hours to ensure they consumed it. As the animals repeated the test at 1 and 3 months, a different flavoured pellet was used each time to prevent bias.

The test was conducted in a cage identical to their home cage over two days. On the first day, the mouse was allowed to explore for 5 minutes the test cage containing only sawdust (3 cm deep). The sawdust from each cage was collected and stored to be used the next day, ensuring the only new variable on the second day was the presence of the pellet. On the second day, the scented pellet was buried in the sawdust, and the animal was given 5 minutes to explore.

#### Hot plate

The hot plate test was employed to evaluate hyperalgesia in mice.^5^ Each mouse was placed on a metallic surface maintained at 52.5°C, surrounded by a 40 cm high plexiglass wall (Digital DS-37 Socrel model; Milan, Italy). The latency to show signs of pain, such as paw licking or jumping, was recorded. If the mouse jumped, it was immediately removed from the plate to prevent injury. A maximum time limit of 45 seconds was set to avoid any tissue damage.

#### Novel object recognition test

The novel object recognition test was conducted to assess cognitive ability. An L-shaped maze arena (8 x 37 x 14.5 cm) was utilized to promote exploration.^6^ The test was performed over three consecutive days, with each day’s session lasting 9 minutes. On the first day (familiarization) the animal was placed in the empty maze. On the second day (acquisition), two identical objects were introduced into the maze, positioned in opposite corners close to the walls. On the third day (test day), one of the identical objects was replaced with a novel object. The animal was placed in the maze facing one of the walls to ensure it did not directly face the objects initially. Exploration and interaction with the objects were monitored and scored using the Behaviour Scoring Panel application (© 2008 by A. DUBREUCQ, version 3.0 beta). The animal was consider to be exploring when it was oriented facing the object and its nose was within a 2 cm radius. Heat map representations of the tests were made with EthoVision® animal tracking system (Noldus, Netherlands). The discrimination index was computed using the formula (Tnew−Told) / (Tnew+Told), where Tnew represents the time spent exploring the novel object, and Told represents the time spent exploring the familiar object. A higher discrimination index indicates a greater preference for the new object over the old one, reflecting successful recognition and memory of the familiar object.

### Statistical analysis

Experimental data were analysed using GraphPad v8.0.1. software (GraphPad Software Inc, U.S.A.), the level of statistical significance was set at p<0.05. All data were tested for outlier identification before statistical analysis and normality was checked using Shapiro-Wilk test with α of 0.05. Data are graphically presented as the group mean ± standard error of the mean (S.E.M.). Graphics depicting only one variable were analysed with Student’s t-test.

All tests were analysed and illustrated comparing two variables: sham or a-syn, and time. These variables were analysed using Two-way ANOVA, with results depicted in the graphics. Post-hoc comparisons using Tukey’s test were shown in the graphics only when the interaction between the two variables (a-syn or time) was significant.

---

**Table 1.** Primary and secondary antibodies used in the immunoassays.

| **Antibody** | **Dilution** | **Reference** | **Purchase from** |
| --- | --- | --- | --- |
| **Primary antibody** |  |  |  |
| Rabbit anti-TH | 1:2000 | AB152 | Sigma-Aldrich |
| Mouse anti-αsyn | 1:10000 | AB27766 | Abcam |
| Mouse anti-αsynP | 1:2000 | AB51253 | Abcam |
| Chicken anti-TH | 1:2000 | AB76442 | Abcam |
| Chicken anti-GFP | 1:400 | AB13970 | Abcam |
| Mouse-anti-NET | 1:1500 | NET05-02 | MAb Technologies |
| **Secondary antibody** |  |  |  |
| Goat anti-chicken Alexa fluor 555 | 1:400 | A21437 | Invitrogen |
| Goat anti-chicken Alexa fluor 488 | 1:400 | AB150169 | Abcam |
| Goat anti-rabbit Alexa fluor 488 | 1:400 | A32731 | Invitrogen |
| Goat anti-rabbit Alexa fluor 555 | 1:400 | A21429 | Invitrogen |
| Donkey anti-mouse Alexa fluor 555 | 1:400 | A31570 | Invitrogen |
| Streptavidine 647 | 1:500 | S21374 | Invitrogen |
| Biotinylated goat anti-rabbit IgG | 1:200 | BA100 | Vector laboratories |
| Biotinylated goat anti-mouse IgG | 1:200 | BA2020 | Vector laboratories |

**Table 2.** Qualitative analysis of human a-syn expression at 1 and 3 months in a unilaterally injected model. The presence of human a-syn-positive axons was evaluated as follows: - no positive axons, + few positive axons, ++ more positive axons, +++ positive axons densely covering the region, ++++ positive cell bodies. IL: Ipsilateral side, injected side. CL: contralateral side.

| Coordinate | General area | Specific area | 1m IL | 1m CL | 3m IL | 3m CL |
| --- | --- | --- | --- | --- | --- | --- |
| $\beta=+1.94$ mm | Prefrontal cortex | | ++ | + | ++ | + |
| $\beta=+0.50$ mm | Striatum | | - | - | + | + |
|  | Accumbens nucleus |  | + | - | + | + |
|  | Lateral septal nucleus |  | + | + | ++ | ++ |
|  | Motor cortex | M1 | ++ | + | ++ | ++ |
|  |  | M2 | ++ | + | ++ | ++ |
| $\beta=+0.38$ mm | Striatum | | - | - | ++ | + |
|  | Lateral septal nucleus |  | + | + | ++ | ++ |
|  | Bed nuclei of the stria terminalis |  | ++ | - | ++ | + |
|  | Motor cortex | M1 | ++ | + | +++ | ++ |
|  |  | M2 | ++ | + | +++ | ++ |
| $\beta=-1.82$ mm | Striatum | | + | + | ++ | + |
|  | Globus pallidus | External section | + | + | ++ | + |
|  | Thalamus | Polymodal association cortex related | + | + | ++ | ++ |
|  | Amygdala | Basolateral | + | + | ++ | + |
|  |  | Central | +++ | ++ | +++ | ++ |
| $\beta=-1.82$ mm | Dorsal hippocampus | CA1 | ++ | ++ | +++ | +++ |
|  |  | CA2 | ++ | ++ | +++ | +++ |
|  |  | CA3 | ++ | ++ | +++ | +++ |
|  |  | DG | ++ | ++ | +++ | +++ |
|  | Thalamus | Polymodal association cortex related | + | + | ++ | ++ |
|  |  | Sensory-motor cortex related | + | + | ++ | ++ |
|  |  | Subthalamic nucleus | +++ | ++ | ++ | ++ |
|  |  | Zona incerta | ++ | + | ++ | ++ |
|  |  | Parasubthalamic nucleus | +++ | + | +++ | ++ |
|  | Amygdala | Basolateral | + | + | ++ | ++ |
|  |  | Central | +++ | ++ | +++ | ++ |
| $\beta=-2.92$ mm | Ventral hippocampus | CA1 | + | + | +++ | ++ |
|  |  | CA2 | + | + | +++ | ++ |
|  |  | CA3 | + | + | +++ | ++ |
|  |  | DG | + | + | +++ | ++ |
|  | Substantia nigra | Pars compacta | ++ | + | ++ | ++ |
|  |  | Pars reticulata | + | + | ++ | ++ |
|  | Ventral tegmental area |  | ++ | ++ | ++ | ++ |
| $\beta=-4.36$ mm | Superior colliculus | Somatosensory | ++ | + | ++ | ++ |
|  |  | Motor | ++ | + | ++ | ++ |
|  | Dorsal raphe nucleus |  | +++ | +++ | +++ | +++ |
| $\beta=-5.40$ mm | Cerebellum | | + | - | ++ | ++ |
|  | Locus coeruleus |  | ++++ | + | ++++ | ++ |
|  | Inferior colliculus |  | ++ | + | ++ | ++ |
|  | Pons | Pontine central gray | +++ | + | ++ | ++ |
|  |  | Parabrachial nucleus | +++ | ++ | +++ | ++ |
|  | Medulla |  | ++ | + | ++ | ++ |

**Table 3.** Action potential properties of sham and a-syn male mice.

|  | **SHAM** | **A-syn** |
| --- | --- | --- |
| **1 month** |  |  |
| Threshold (mV) | -56.79 ± 2.33 | -57.91 ± 2.79 |
| AP amplitude (mV) | 80.92 ± 2.38 | 81.25 ± 1.27 |
| AHP amplitude (mV) | -23.45 ± 0.76 | -25.46 ± 0.77 |
| Half-width (ms) | 1.04 ± 0.06 | 0.97 ± 0.03 |
| **3 months** |  |  |
| Threshold (mV) | -51.43 ± 3.50 | -58.31 ± 2.40 |
| AP amplitude (mV) | 75.46 ± 3.39 | 77.44 ± 2.49 |
| AHP amplitude (mV) | -20.06 ± 0.64 | -20.03 ± 1.10 |
| Half-width (ms) | 1.35 ± 0.05 | 1.30 ± 0.09 |

Data are expressed as the mean ± S.E.M. of n cell.

**Table 4.** Action potential properties of sham and a-syn female mice.

|  | **SHAM** | **A-syn** |
| --- | --- | --- |
| **1 month** |  |  |
| Threshold (mV) | -50.44 ± 2.51 | -61.10 ± 2.68* |
| AP amplitude (mV) | 77.73 ± 1.85 | 80.99 ± 2.28 |
| AHP amplitude (mV) | -22.58 ± 0.58 | -24.16 ± 1.01 |
| Half-width (ms) | 0.88 ± 0.06 | 0.75 ± 0.03 |
| **3 months** |  |  |
| Threshold (mV) | -59.94 ± 2.84 | -59.08 ± 2.97 |
| AP amplitude (mV) | 76.71 ± 2.06 | 83.73 ± 2.04 |
| AHP amplitude (mV) | -21.94 ± 1.04 | -21.25 ± 0.87 |
| Half-width (ms) | 1.39 ± 0.09 | 1.21 ± 0.03 |

Data are expressed as the mean ± S.E.M. of n cell.
\* $P < 0.05$ . Student's t-test

